# More Signal, Less Noise: Single-primer Adenylation Method (SAM) for the Efficient Preparation of DNA for Transposon Sequencing (TnSeq)

**DOI:** 10.1101/012732

**Authors:** S Willcocks, R Stabler, B Wren

## Abstract

Transposon Sequencing (TnSeq) is a cutting-edge tool that allows quantitative, genome-wide genetic analysis of a given population in an experimental condition. TnSeq utilises next generation sequencing that is designed to perform optimally when all of the input material is of interest, therefore TnSeq is inefficient since only a small portion of nucleic acid fragments contain the transposon target site for primer annealing. We describe an innovative modification to standard transposon library preparation protocols that provides a significant improvement to the quality of sequencing data recovered.

## INTRODUCTION

The advent of next generation sequencing is revolutionising infectious disease research and holds great promise, particularly in elucidating the evolution and spread of organisms and in clinical diagnostics (Alfoldi and Lindblad-Toh 2013). When coupled with the creation of genome-wide transposon-mutant libraries, transposon-sequencing (TnSeq) has emerged as a powerful tool that allows the simultaneous assessment of every gene in a given genome, revealing essentiality in the chosen model - be this antibiotic resistance, virulence, or simply survival in optimal growth media (van Opijnen, Bodi et al. 2009; Gallagher, Shendure et al. 2011; Moule, Hemsley et al. 2014). TnSeq requires first the generation of a transposon-mediated mutant library that saturates the genome, so that every gene carries multiple insertions; individual mutants are grown on agarose plates and then collected and pooled prior to experimentation. Sequencing of the unknown region of genomic (g)DNA flanking the transposon reveals which gene has been mutated. Comparison of the genes represented in the ‘output’ versus the ‘input’ library reveals, in a quantitative manner, negative selection events *ie* mutation of genes that are required for survival in the test condition precludes their detection in the output pool (Chaudhuri, Morgan et al. 2013). The applications of TnSeq are vast; as with all new technologies, however, there are technical hurdles.

## RESULTS AND DISCUSSION

The widely used protocol for preparing DNA for TnSeq, as with Illumina related platforms, involves the fragmentation of extracted DNA, ligation of adaptor sequences, and the selective amplification of transposon-containing fragments (van Opijnen, Lazinski et al. 2014). What separates TnSeq from standard sequencing of gDNA or RNA-derived extracts, is that only a very small percentage of the genetic material contains the target of interest: the transposon (**Fig 1A**). This radically reduces the efficiency and statistical power of next generation sequencing. A common solution is to amplify, by means of polymerase chain reaction (PCR) with specific oligos, only fragments of DNA that contain the transposon. PCR is crucial to TraDIS since it is also used to introduce the sequence that allows the DNA fragments to anneal to the surface of the flow cell. While PCR improves the signal to noise ratio of target to background DNA, it is also the main step through which bias is introduced into the data (Kozarewa, Ning et al. 2009; Aird, Ross et al. 2011). Bias seems to be exacerbated in low-complexity (hundreds-scale pools) libraries, and somewhat masked in very large (hundreds of thousands scale) libraries.

Limiting amplification bias through primer design and changes to cycle parameters such as melting time and ramp speed may provide modest improvements to PCR efficiency. However, since the target site for primer annealing is conserved regardless of organism or location of the transposon, this is not a satisfactory candidate for explaining PCR amplification bias. More crucial are the characteristics of the nucleic acids outside the region of the transposon including sequence, fragment length, and GC content, that can all adversely affect melting temperature and primer annealing efficiency through secondary structure formation (Day, Speiser et al. 1996; Barnard, Futo et al. 1998; Ogino and Wilson 2002; Dohm, Lottaz et al. 2008). For this reason, TnSeq of organisms with unusually high (*Burkholderia spp*) or low (*Yersinia spp*) GC content, or even those with relatively balanced GC content, but with interspersed regions of high GC content (*Escherichia coli*), can be problematic. Bias in amplification can result in apparent ‘hotspots’ that are not explained by the natural favouring by the transposon of particular conserved sequences. Worse, it can result in false conclusions of positivity or negativity.

It has been noted by other researchers, that lowering the threshold of PCR to no more than 10 cycles allows enrichment of the transposon-containing fragments, while limiting the bias to an acceptable level. However, this exacerbates the problem of low signal to background with TnSeq. The user is required to decide the optimal DNA concentration to apply to the flow cell, which requires a narrow range for optimal results. One either normalises DNA concentration to the amount of total DNA, in which case the percentage of clusters that contain transposon sequence is very small, or one normalises to the concentration of transposon-only DNA, in which case the non-transposon ‘background’ DNA is so high that it prevents capture of good quality images by the sequencing machine software. This desultory element in terms of choosing PCR cycle number or DNA concentration is prohibitive; it is expensive in time and money and more importantly in precious samples that may not readily be re-attainable.

One solution to the issue of high background is to attempt to recover or enrich the transposon-positive fragments post-PCR. We have investigated methods including the enzymatic digest of non-PCR DNA, theoretically leaving only the transposon-positive amplicons, as well as the use of biotinylated primers and nano-bead capture technology to physically separate the target DNA of interest. However, with both of these methods, we found a loss of yield and no significant enhancement of signal to background as assessed by quantitative PCR.

We presently report an innovation that allows for the exclusive binding of transposon-containing DNA fragments to the flow cell, resulting in clean, high quality data that is highly reproducible. The use of the Single-primer Adenylation Method (SAM) for TnSeq obviates the need to find an elusive balance between having enough cycles to amplify the transposons whilst keeping bias to a minimum. We are able to routinely use just 10 cycles of PCR, even on low complexity libraries to prepare a library with very limited bias, with negligible background from non-transposon DNA. Essentially, more signal and less noise.

In order for any DNA to bind to the Illumina MiSeq flow cell surface during a sequencing run, it must contain at least one of two defined sequences, ‘P5’ and ‘P7’; these are complementary to a lawn of primers that are pre-coated onto the flow cell. The P5 and P7 sequences are introduced by PCR that uses adapter sequences as a template for reverse strand synthesis, to introduce P7, and the transposon serves as the template for the forward primer, to introduce P5. Since PCR amplification with this primer pair only occurs with transposon-containing fragments, one may expect that only transposon-containing fragments have the P5 and P7 sequence introduced and therefore only transposon fragment are able to bind to the flow cell.

However, non-transposon DNA fragments are present in much higher amounts than transposon-containing fragments. It is therefore possible that, although no amplification takes place, a single extension of complementary DNA that carries the P7 sequence on the reverse primer would occur with the large proportion of non-transposon DNA that would subsequently be capable of binding to the flow cell. This portion of DNA would not be detected by qPCR methods that use either P5 or transposon sequence for the forward primer.

The use of adapters is essential, since it serves as a template for the reverse primer during amplification of transposon-containing fragments. The ligation process depends on the indiscriminate adenylation of the 3′ end of all DNA fragments using an enzyme such as Klenow fragment, exo-. Alternatively, by enzymatic digest of the transposon to leave an overhang, the efficiency of which depends on the frequency of the chosen restriction sequence in the target genome and usually, sequence modification of the transposon itself (van Opijnen, Lazinski et al. 2014). This is the key step that we have revised, so that only transposon-containing fragments receive the A-tail, forbidding the generation of P7-tagged non-transposon fragments and with no need for restriction digest. In our revised protocol: genomic DNA extraction, fragmentation and end repair still occur as per standard preparation protocols. The critical step is to conduct a PCR using a single transposon-specific primer to introduce a 3′ A-overhang using *Taq* polymerase, instead of adenylating all fragments using Klenow fragment, exo-. The forward primer ideally is identical to the transposon-specific primer carrying the P5 tag that is used in the subsequent amplification PCR (**Fig 1B**).

The need for high fidelity is clear, and there are numerous products available commercially that contain a blend of a high fidelity polymerase with a standard, A-tailing *Taq* polymerase and the PCR parameters are as per manufacturer’s recommendations for the chosen polymerase. Since there is no amplification during single-primer adenlyation, multiple cycles may be used without concern of bias introduction. Additives to assist with GC melting and an extended melting cycle time can also be used effectively at the discretion of the user, depending on the requirements of the source material. Additionally, there is the potential for size selection during single-primer adenylation: by controlling the extension time, one can dictate the size of fragments that are A-tailed. We recommend either a slow ramp-speed or a final slow cool-down step to assist correct annealing of the A-tailed strand with its complementary template.

After single-primer adenylation, the result is that the only location for the adapters to ligate to, are the ends of the transposon-positive fragments. Our data suggest that at least 50% of transposon-positive fragments are A-tailed this way.

The samples are then ready for amplification PCR as per standard TnSeq protocols, using the P5-containing transposon-specific forward primer, and the P7-containing adapter-specific reverse primer. Splitting the input material into several independent reaction tubes to be pooled post-PCR is recommended to reduce PCR amplification bias, as is a maximum of 10 cycles. Adapter-ligated transposon-containing fragments may be quantified using P5 and P7 specific primers for quantitative PCR prior to sequencing using Illumina MiSeq.

SAM for TnSeq in contrast with existing methodology for DNA preparation is summarised in Fig 1:

**Fig 1.**
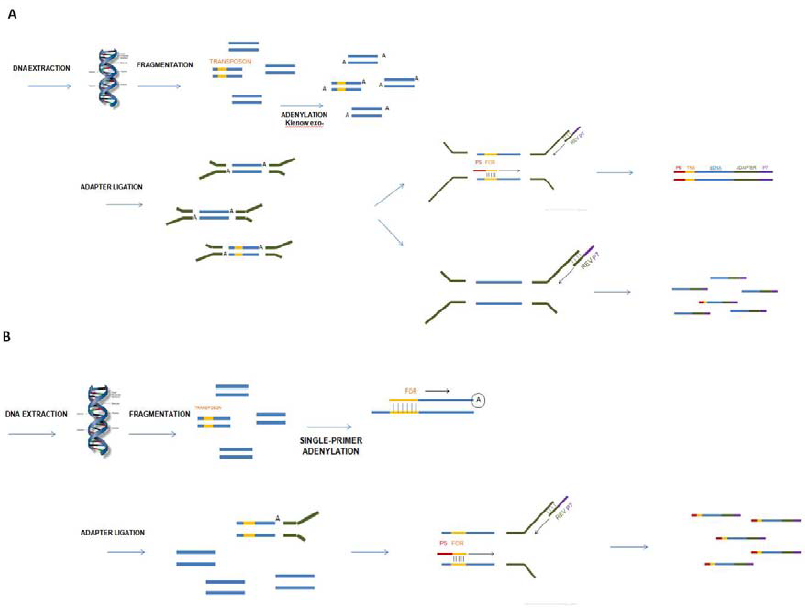
Single-primer Adenylation Method (SAM) for TnSeq. In the standard DNA preparation protocol for TnSeq (**A**), DNA fragments (blue) are indiscriminately adenylated, resulting in all fragments ligating with adapters (green) regardless of whether they contain the transposon (yellow). Even in the absence of transposon DNA, the reverse primer that introduces the P7 flow-cell binding sequence (purple) may still be introduced in the majority of fragments, although they are not subsequently amplified by PCR. This reduces the signal:background ratio and reduces efficiency of TnSeq. Using the Single-primer Adenylation Method (SAM) (**B**), a transposon-specific primer is used to selectively introduce an A-tail only to transposon-containing fragments. Subsequently, only these fragments are able to ligate with adapters, which may then be amplified by PCR in the usual manner.

Only transposon-positive fragments are amplified and able to bind to the flow cell surface, virtually eradicating background signal from the majority of DNA that does not contain transposon DNA. It is possible that some clusters are still generated that seemingly do not yield useful transposon location information. This may be due to non-specific annealing of fragments to the flow-cell primers, which by definition are able to form clusters since the ‘Read One’ primer used by Illumina is related to the flow cell primer sequence. Alternatively, clusters may indeed contain transposon-containing sequences that are missed from post-sequencing analysis algorithms due to mismatches, and this will depend on the stringency demands set by the user.

The efficiency of SAM for TnSeq is enhanced over standard preparation protocols in two respects. First, only transposon-positive fragments are able to be ligated to the adapter sequence to serve as a template for amplification PCR. Second, only these fragments are able to receive the P5 sequence required for maximal binding to the flow cell, as this is included in the forward primer during the amplification step. With our method, non-transposon DNA fragments possess neither P7 nor P5 sequences, whereas in standard protocols, they contain both. This results in less total DNA binding to the flow-cell, and less cluster formation, but significantly more of these clusters map to the transposon-containing regions of DNA. Measurement of total DNA by methods such as Bioanalyzer or Nanodrop is of limited value in SAM for TnSeq, since this quantifies DNA that is not capable of binding to the flow-cell. Therefore qPCR using transposon-specific primers is essential to adjust the DNA to the appropriate concentration. Using SAM for TnSeq, quantification that uses a P7-specific primer is more accurate than the standard protocol, since P7 is only carried by transposon-containing fragments.

In conclusion, our method greatly enhances the efficiency of TnSeq, by virtue of a simple modification to an early step in the preparation of DNA for sequencing. The revised method is applicable for all TraDIS experiments, and is of particular value for researchers with challenging, GC-polarised genomic material, and those with small, low complexity libraries.

## MATERIALS AND METHODS

Genomic DNA is extracted from the experimental organism followed by fragmentation by either sonic disruption or a frequently-cutting restriction enzyme and then purified. The ends of the DNA fragment are repaired to make blunt ends using (for a 75 μl sample): 10 μl phosphorylation buffer, 4 μl dNTPs, 1.5 μl Klenow DNA polymerase, 5 μl T4 PNK, 5 μl T4 DNA polymerase at room temperature (RT) for 30 mins. The reaction is purified and then subjected to single-primer adenlyation. A blended polymerase containing A-tailing *taq* and a proof-reading enzyme is recommended, such as Advantage Polymerase (Takara Clontech) and the PCR parameters are as per manufacturer’s recommendations for the chosen polymerase. The forward primer ideally is identical to the transposon-specific primer carrying the P5 tag that is used in the subsequent amplification PCR (using a high fidelity enzyme such as NEBNext or Phusion). Since there is no amplification during single-primer adenlyation, multiple cycles may be used without concern of bias introduction. If desired, the extension time may be intentionally restricted to limit the size of fragments that are A-tailed. We recommend either a slow ramp-speed or a final slow cool-down step to assist correct annealing of the A-tailed strand with its complementary template. Following PCR purification, DNA fragments are ligated with annealed adapters according to standard TnSeq procedure and again purified. The samples are then ready for amplification PCR, using the P5-containing transposon-specific forward primer, and the P7-containing adapter-specific reverse primer. Splitting the input material into several independent reaction tubes to be pooled post-PCR is recommended to reduce PCR amplification bias, as is a maximum of 10 cycles. Adapter-ligated transposon-containing fragments may be quantified using P5 and P7 specific primers for quantitative PCR prior to sequencing using Illumina MiSeq.

PCR parameters for Single-Primer Adenylation Using Advantage Polymerase:

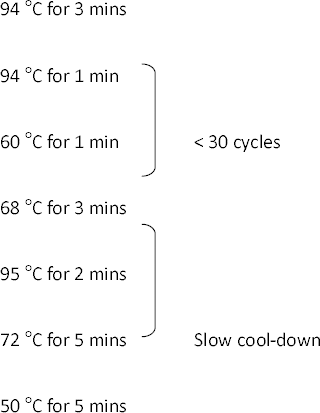

